# GC-biased gene conversion in X-Chromosome palindromes conserved in human, chimpanzee, and rhesus macaque

**DOI:** 10.1101/2021.04.21.440828

**Authors:** Emily K. Jackson, Daniel W. Bellott, Helen Skaletsky, David C. Page

## Abstract

Gene conversion is GC-biased across a wide range of taxa. Large palindromes on mammalian sex chromosomes undergo frequent gene conversion that maintains arm-to-arm sequence identity greater than 99%, which may increase their susceptibility to the effects of GC-biased gene conversion. Here, we demonstrate a striking history of GC-biased gene conversion in 12 palindromes conserved on the X chromosomes of human, chimpanzee, and rhesus macaque. Primate X-chromosome palindrome arms have significantly higher GC content than flanking single-copy sequences. Nucleotide replacements that occurred in human and chimpanzee palindrome arms over the past 7 million years are one-and-a-half times as GC-rich than the ancestral bases they replaced. Using simulations, we show that our observed pattern of nucleotide replacements is consistent with GC-biased gene conversion with a magnitude of 70%, similar to previously reported values based on analyses of human meioses. However, GC-biased gene conversion explains only a fraction of the observed difference in GC content between palindrome arms and flanking sequence, suggesting that additional factors are required to explain elevated GC content in palindrome arms. This work supports a greater than 2:1 preference for GC bases over AT bases during gene conversion, and demonstrates that the evolution and composition of mammalian sex chromosome palindromes is strongly influenced by GC-biased gene conversion.

## INTRODUCTION

Homologous recombination maintains genome integrity through the repair of double-stranded DNA breaks, while also promoting genetic innovation through programmed reshuffling during meiosis. Homologous recombination can produce crossover events, in which genetic material is exchanged between two DNA molecules, or non-crossover events. Crossover events and non-crossover events both result in gene conversion, the non-reciprocal transfer of DNA sequence from one homologous template to another. When the templates involved in gene conversion are not identical, gene conversion can be biased, resulting in the preferential transmission of one allele over another (reviewed in Galtier et al. 2001, Marais 2003, Duret and Galtier 2009). In particular, GC alleles are generally favored over AT alleles, leading to a strong correlation between GC content and recombination rates across the genome. GC-biased gene conversion is widespread across taxa, including plants (Muyle et al. 2011), yeast (Mancera et al. 2008), birds (Smeds et al. 2016), rodents (Montoya-Burgos et al. 2003, Clément and Arndt 2011), humans (Odenthal-Hesse et al. 2014, Williams et al. 2015, Halldorsson et al. 2016), and other primates (Galtier et al. 2009, Borges et al. 2019).

While early evidence for GC-biased gene conversion was indirect (Galtier et al. 2001, Marais 2003), two recent studies identified gene conversion events in humans directly using three-generation pedigrees (Williams et al. 2015, Halldorsson et al. 2016). This approach enabled calculation of the magnitude of GC bias, defined as the frequency at which gene conversion at a locus containing one GC allele and one AT allele results in transmission of the GC allele. Williams et al. identified 98 autosomal non-crossover gene conversion events at loci with one GC allele and one AT allele, and found that 63 (68%) transmitted the GC allele (Williams et al. 2015). Halldorsson et al. analyzed autosomal crossover and non-crossover gene conversion events separately, and found GC biases of 70.1% and 67.6%, respectively (Halldorsson et al. 2016). The magnitude of GC bias may vary across different genomic positions: Another study used sperm typing to examine allele transmission at six autosomal recombination hotspots, and found evidence for GC-biased transmission at two hotspots, but unbiased transmission at the other four hotspots (Odenthal-Hesse et al. 2014).

Mammalian sex chromosomes contain large, highly identical palindromes, with arms that can exceed 1 Mb in length and arm-to-arm identities greater than 99% (Skaletsky et al. 2003, Warburton et al. 2004, Hughes et al. 2010, Mueller et al. 2013, Soh et al. 2014, Hughes et al. 2020, Jackson et al. 2020). Near-perfect identity between palindrome arms is maintained by high rates of ongoing gene conversion (Rozen et al. 2003), which may make palindromes uniquely susceptible to the effects of GC-biased gene conversion (Hallast et al. 2013, Skov et al. 2017). Recently, we generated high-quality reference sequence for twelve large palindromes that are conserved on the X chromosomes of human, chimpanzee, and rhesus macaque, demonstrating a common origin at least 25 million years ago (Jackson et al. 2020). Here, we use a comparative genomic approach combined with evolutionary simulations to analyze the impact and magnitude of GC-biased gene conversion in primate X-chromosome palindromes. We find that GC content is elevated in palindrome arms relative to flanking sequence, and that recent nucleotide replacements in human and chimpanzee palindrome arms are approximately one-and-a-half times as GC-rich as the ancestral bases that they replace. Using simulations of palindrome evolution, we show that our observed pattern of nucleotide replacements is consistent with a magnitude of GC bias of about 70%, which supports recent estimates derived from analyses of human meioses using an orthogonal approach.

## MATERIALS AND METHODS

### Human mutation rate

Three recent publications used whole-genome shotgun sequencing data from related individuals to calculate human mutation rates of around 1.2×10^−8^ mutations per nucleotide per generation (Roach et al. 2010, Kong et al. 2012, Jónsson et al. 2017). However, these publications used only autosomal data, while the human X chromosome may have a lower mutation rate than autosomes due to its unique evolutionary history (Schaffner 2004). To our knowledge, similarly high-quality estimates of the human X chromosome mutation rate do not exist. To estimate the mutation rate for the human X chromosome, we examined Supplemental Table 4 from Jónsson et al., which provides information for all autosomal and X chromosome mutations detected in their dataset. Supplemental Table 4 reports 2694 X chromosome mutations from 871 probands, or around 3.1 mutations per generation. To calculate the autosomal mutation rate, Jónsson et al divided the number of autosomal mutations per generation by the number of autosomal base pairs with adequate coverage depth in their dataset. We therefore divided 3.1 X-chromosome mutations per generation by the length of the X chromosome in hg38 (156,040,895 base pairs) multiplied by the fraction autosomal coverage (93.3%), which we assume here is similar to the fraction of X chromosome coverage. This approach yielded an estimated human X chromosome mutation rate of 1.06 ×10^−8^ mutations per nucleotide per generation. This value is about 20% lower than the value calculated by Jónsson et al. for autosomes (1.28 × 10^−8^ mutations per nucleotide per generation), consistent with predictions that mutation rates are lower on X chromosomes than on autosomes.

### GC content of primate X-chromosome palindromes

We calculated the GC content for each palindrome (Arm 1, spacer, and flanking sequence) using custom Python code. We performed all analyses using clones sequenced by Jackson et al. 2020. For flanking sequence, we used available sequence upstream and downstream of palindrome arms that was present in all three species. For example, if the human clones for a given palindrome contained 3’ sequence that was not sequenced in chimpanzee and rhesus macaque, we trimmed the human sequence to contain only the portion alignable between all three species. Visualizations were generated using ggplot2 in R (Wickham 2016, R Core Team 2020).

### Generation of sequence alignments

Sequence alignments were performed using ClustalW with default parameters (Thompson et al. 1994). To identify and exclude regions of poor alignment, ClustalW sequence alignments were scanned using a sliding 100-bp window and filtered to exclude windows with fewer than 60 matches between species, using custom Python code (Jackson et al. 2020).

### Calculation of divergence

Divergence was calculated by generating pairwise alignments using ClustalW, then calculating p-distance with MEGA X (Kumar et al. 2018). For alignment of arms between species, we generated pairwise alignments using Arm 1 from each species (Jackson et al. 2020).

### Simulations

Our simulations were designed to model the evolution of a palindrome present in the common ancestor of human, chimpanzee, and rhesus macaque, and maintained in all three lineages until the present. For each iteration, we initialized an ancestral palindrome with each nucleotide chosen at random based on the median characteristics of conserved primate X-chromosome palindromes (arm length: 37 kb, arm-to-arm identity: 99.953%, GC content: 46%). Each ancestral palindrome then underwent rounds of substitution followed by intra-chromosomal gene conversion, with two branching events to account for the divergence of human, chimpanzee, and rhesus macaque (see below for the calculation of the number of generations in each branch). Simulation parameters included the substitution rate for each evolutionary branch, relative rates for different types of substitutions (i.e., the neutral substitution matrix), and the frequency and GC bias of intra-chromosomal gene conversion, with parameter values selected as described below. Simulations were implemented with custom Python code.

### Estimation of generation numbers for simulations

Divergence times for human versus chimpanzee and for human versus rhesus macaque are estimated at about 7 and 29 million years, respectively (Kumar et al. 2017). Generation times for primates vary between species, with estimated generation times around 30 years for humans (Tremblay and Vézina 2000, Matsumura and Forster 2008), 25 years for chimpanzee (Langergraber et al. 2012), and 11 years for rhesus macaque (Gage 1998, Xue et al. 2016). For simplicity, we assumed an intermediate value of 20 years per generation for all branches. Using these values, we estimated a total of 1,450,000 generations for the branch from the common human-chimpanzee-rhesus macaque (HCR) ancestor to rhesus macaque (Branch 1), 1,100,000 generations for the branch from the common HCR ancestor to the common human-chimpanzee (HC) ancestor (Branch 2), and 350,000 generations each for the branches from the common HC ancestor to chimpanzee and to human (Branches 3 and 4, respectively). For a discussion of the impact of generation numbers on our simulations, see Supplemental Note 2.

### Estimation of substitution rates for simulations

Substitution rates per generation can be inferred from the nucleotide divergence observed between species of known divergence times. We calculated these rates for each branch of our simulated evolutionary tree as follows:

> *Substitution rate: Human versus chimpanzee*

> Palindrome arm divergence: 0.84% (Jackson et al. 2020)
>
> Generations: 350,000 * 2 = 700,000 (see above)
>
> Substitution rate: 1.20 × 10^−8^ substitutions per base per generation.

> *Substitution rate: Human versus rhesus macaque*

> Palindrome arm divergence: 5.4% (Jackson et al. 2020)
>
> Generations: 1,450,000 * 2 = 2,900,000 (see above)
>
> Substitution rate: 1.86 × 10^−8^ substitutions per base per generation.

The human-chimpanzee substitution rate was mapped directly onto Branches 3 and 4. The human-rhesus macaque substitution rate was mapped directly onto Branch 1. For Branch 2, we calculated the substitution rate such that the expected divergence along Branch 1 would equal the expected divergence along Branch 2 + Branch 3:

> 2.7% = 0.42% + (Branch 2 rate * 1,100,000 generations)
>
> Branch 2 rate: 2.07 × 10^−8^ substitutions per base per generation.

Note that for the Branch 2 calculation we assume symmetry of divergence, i.e., divergence between two lineages is divided equally between them.

To confirm that our substitution rates were reasonable, we converted our values to per-year mutation rates assuming a generation time of 20 years, and compared these rates to previously published values. All three of our per-year substitution rates fall within confidence intervals for the same species estimated using autosomal data (Scally and Durbin 2012). Our values fell near the lower end of the confidence intervals, consistent with the prediction that substitution rates on the X chromosome should be slightly lower than on autosomes. Note that our estimated substitution rates per generation differ from the mutation rates reported above for the human X chromosome: Single-generation mutation rates are known to differ from substitution rates over long evolutionary timescales, likely due to a recent slowdown in the mutation rate in humans and great apes (Scally et al. 2012). For a discussion of the impact of mutation rates on our simulations, see Supplemental Note 2.

### Estimation of neutral substitution matrix for simulations

Neutral substitution patterns between species do not follow a uniform distribution: Transitions are more common than transversions, and substitutions that replace a strong base (GC) with a weak base (AT) are more common than substitutions in the opposite direction (Petrov and Hartl 1999, Zhang and Gerstein 2003, Duret and Arndt 2008). In addition to branch-specific substitution rates, we therefore also sought to determine a reasonable pattern of neutral substitutions for our simulations.

We identified neutral substitutions using alignments from 3.8 Mb of gene-masked sequence flanking X-chromosome palindromes, using parsimony to infer substitution events in human and chimpanzee with rhesus macaque as an outgroup. From this we calculated seven different substitution rates:

| Substitution | Substitution rate (substitutions/nt/ generation) |
| --- | --- |
| AT →TA | $1.64 \times 10^{-9}$ |
| AT →CG | $1.93 \times 10^{-9}$ |
| AT →GC | $8.04 \times 10^{-9}$ |
| CG → GC | $2.98 \times 10^{-9}$ |
| CG →AT | $3.22 \times 10^{-9}$ |
| CG → TA (non-CpG) | $1.02 \times 10^{-8}$ |
| CG→TA (CpG) | $9.58 \times 10^{-8}$ |

The overall neutral substitution rate (K) can be calculated as described in Duret and Arndt 2008:

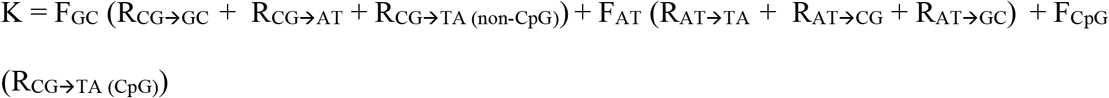

where F_GC_, F_AT_ and F_CpG_ represent the frequencies of each site and R_AA_→_BB_ represents the frequencies of each substitution. Using the substitution rates above combined with the observed frequencies of each site (F_GC_: 0.396, F_AT_: 0.596, F_CpG_: 0.08), we found that K = 1.42 × 10^−8^ substitutions per nucleotide per generation. We then combined the categories CG→TA (non-CpG) and CG→TA (CpG) into a single rate CG→TA as follows:

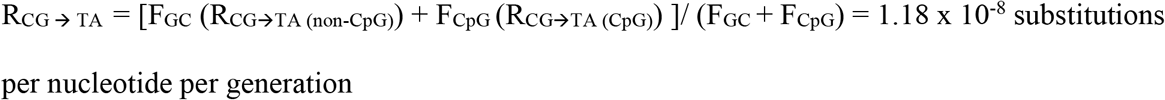

We do not expect combining rates for CpG and non-CpG substitutions to affect either of our simulation output metrics (Figures 3B-C: Fraction GC derived – Fraction GC ancestral at sites of nucleotide replacements; Figure 3D: Fraction GC overall) because these metrics are agnostic to the context in which each fixed nucleotide replacement occurred.

**Figure 1.**
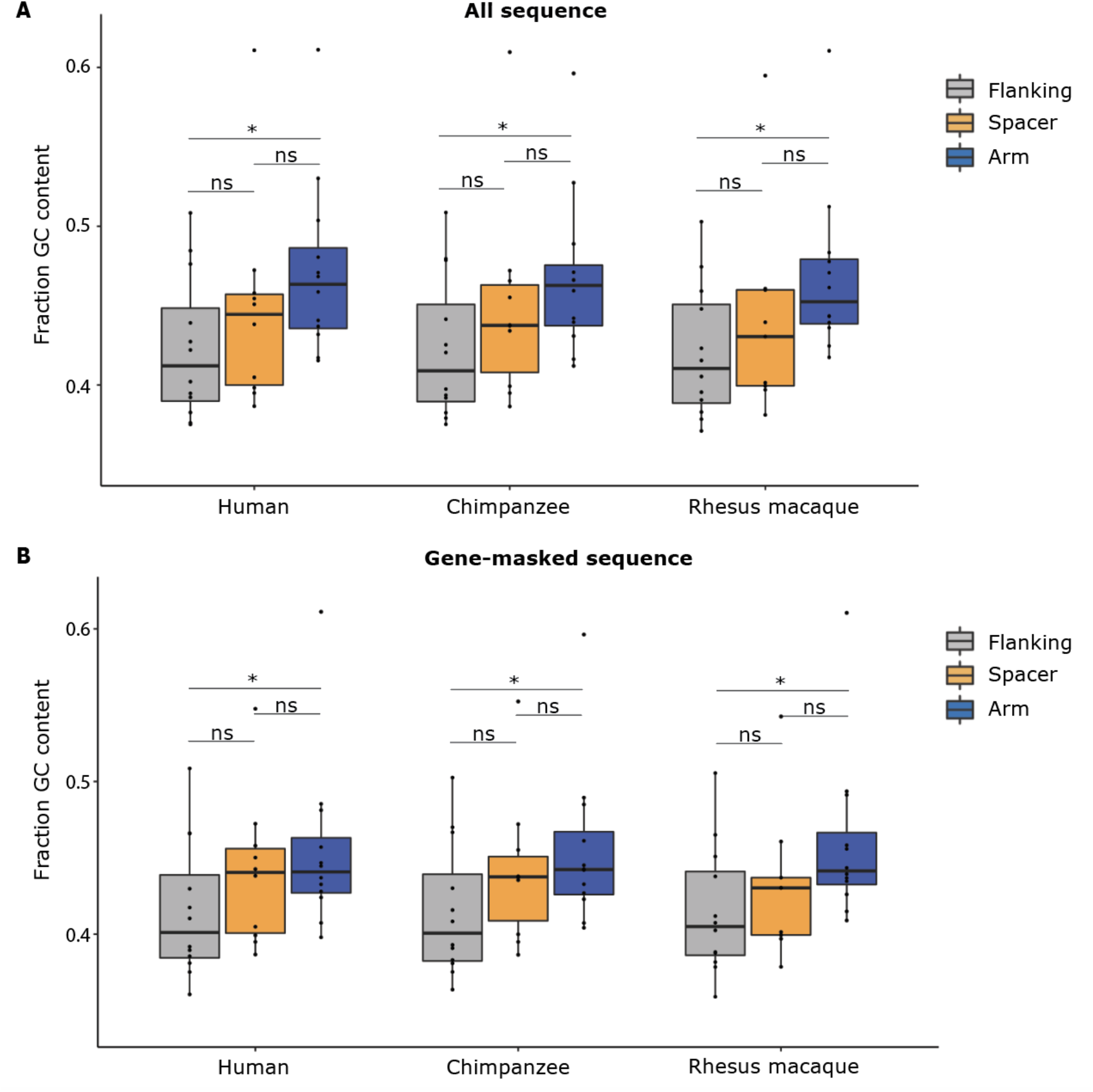
GC content is elevated in primate X-chromosome palindrome arms compared to flanking sequence. GC content measured in 12 palindromes conserved between human, chimpanzee, and rhesus macaque. Small spacers (<5 kb) excluded from analysis. Results a) for all sequence and b) after masking protein-coding genes (gene body plus 1 kb upstream). *p<0.05, ns = not significant, Mann-Whitney U.

**Figure 2:**
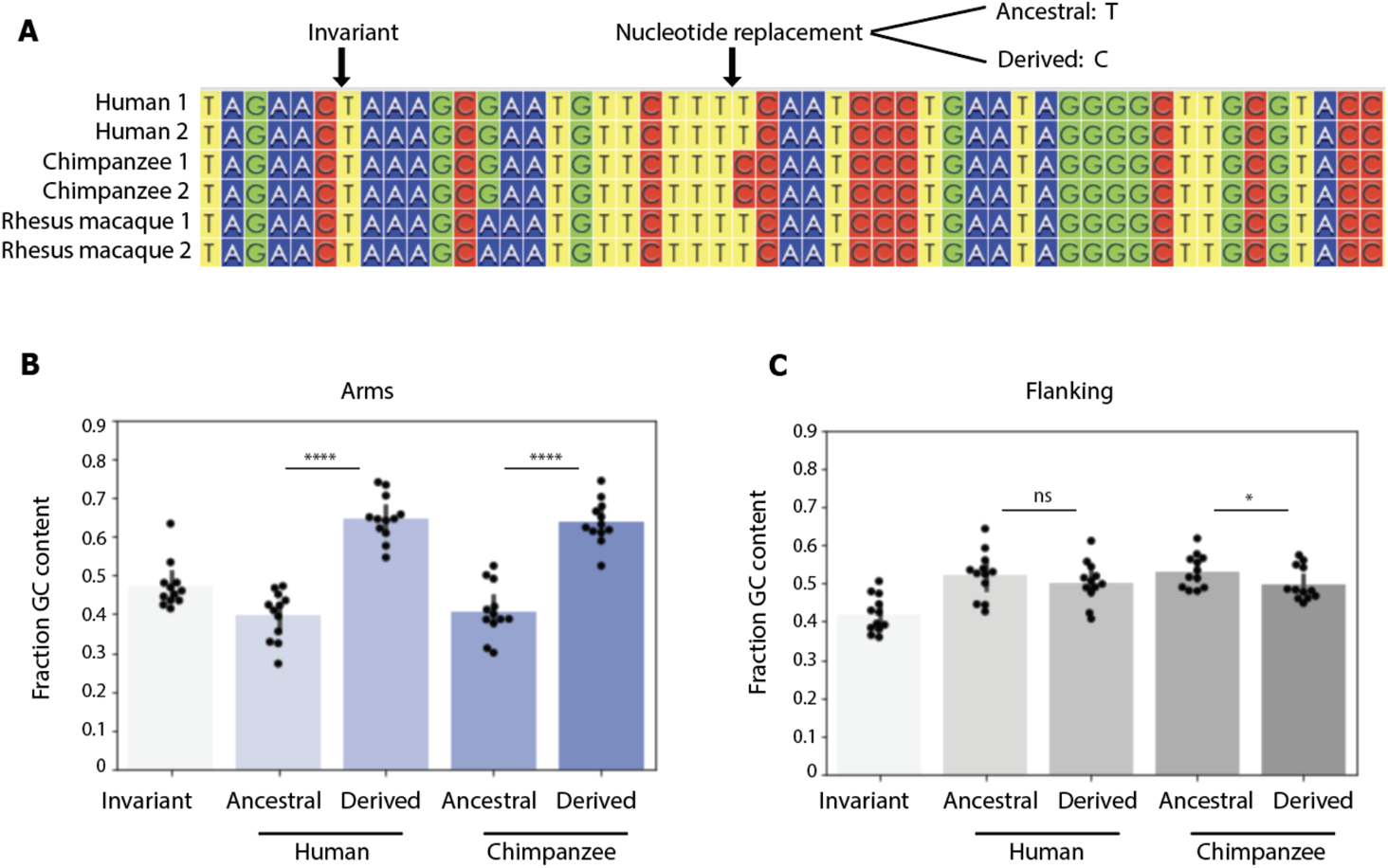
Nucleotide replacements in human and chimpanzee X-chromosome palindrome arms in the past 7 million years have been GC-biased. a) Identification of nucleotide replacements from six-way arm alignments from palindromes conserved between human, chimpanzee, and rhesus macaque. Invariant sites are identical in human, chimpanzee, and rhesus macaque. Alignments generated with ClustalW and visualized using Wasabi (Veidenberg et al. 2016). b,c) Fraction GC content for ancestral versus derived bases. ****p<0.0001, *p<0.05, Mann-Whitney U.

**Figure 3:**
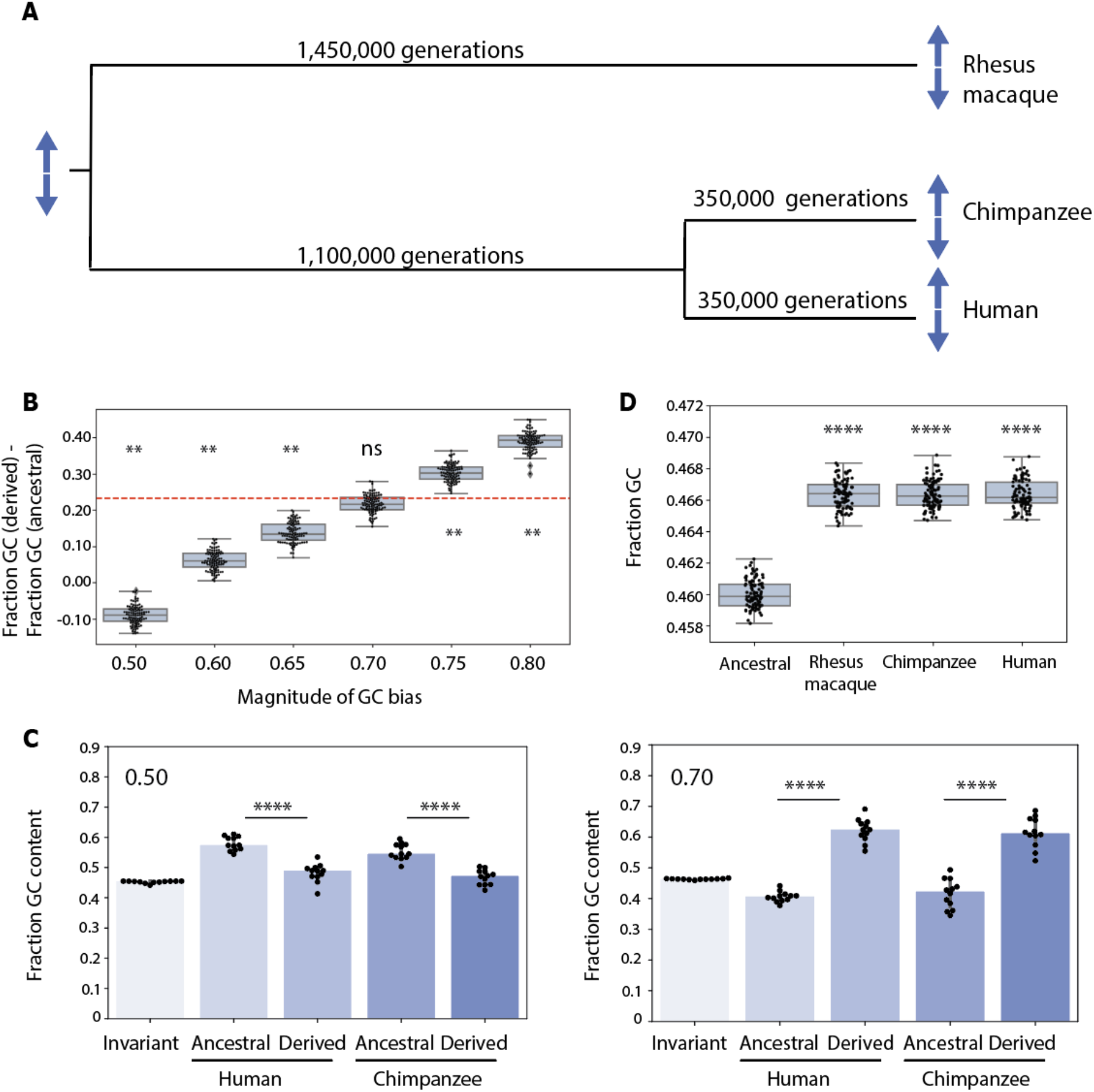
Simulating palindrome evolution with different degrees of GC bias. a) Schematic of simulations. b) Simulated differences between GC content of ancestral and derived bases for six different magnitudes of GC bias. Each dot (n=100 for each magnitude of GC bias) represents the median difference for a set of 12 simulated palindromes. Dashed red line represents true value observed in Figure 2B. **p<0.01, ns = not significant, bootstrapping. c) Fraction GC content for ancestral versus derived bases in simulated palindromes. Results shown for one representative set of 12 palindromes from simulations in Figure 3B. Upper left corner: Magnitude of GC bias. ****p<0.0001, *p<0.05, Mann-Whitney U. d) Fraction GC content for simulated palindrome arms and ancestral sequence. Magnitude of GC bias = 0.70. Each dot (n=100 for each category) represents median GC content for a set of 12 simulated palindromes. ****p<0.0001, *p<0.05, Mann-Whitney U.

The substitution rates above were calculated using substitutions in flanking sequence since the divergence of chimpanzee and human; however, each evolutionary branch in our simulation has a different overall substitution rate (see section above). For each branch, we therefore divided the substitution rates above by the original overall substitution rate of 1.42 × 10^−8^ substitutions per nucleotide per generation, then multiplied by the branch-specific overall substitution rate. This kept the relative ratios between different substitution types constant, while accounting for different overall substitution rates in each branch. The effects of reasonable alterations of this neutral substitution matrix, including adjusting for possible under-estimation of the CpG substitution rate due to artifacts of parsimony, are described in Supplemental Note 3.

## RESULTS

### High rates of intrachromosomal gene conversion in arms of primate X-chromosome palindromes

To understand the role of GC-biased gene conversion in the evolution of primate X-chromosome palindromes, we first calculated the rate of intrachromosomal gene conversion between palindrome arms. Sequence identity between palindrome arms depends on the balance between two evolutionary forces: The rate at which new mutations arise in each arm, and the rate at which gene conversion between arms homogenizes the resulting sequence differences. The rate of intrachromosomal gene conversion can therefore be calculated using the formula c = 2μ /d, where μ represents the mutation rate, and d represents the fraction divergence between arms (Rozen et al. 2003). Among twelve X-chromosome palindromes conserved between human, chimpanzee, and rhesus macaque, we found a median divergence between arms of 4.7 × 10^−4^ differences per nucleotide, or around one difference per 2200 nucleotides. Assuming a mutation rate of 1.06 × 10^−8^ mutations per nucleotide per generation (Roach et al. 2010, Kong et al. 2012, Jónsson et al. 2017, see Methods), we calculated a gene conversion rate of 4.5 × 10^−5^ events per nucleotide per generation for primate X-chromosome palindromes. This value is nearly eight times the recent estimate of 5.9 × 10^−6^ gene conversion events per nucleotide per generation across human autosomes (Williams et al. 2015, Halldorsson et al. 2016), highlighting the rapid pace of genetic exchange between sex chromosome palindrome arms.

### GC content is elevated in primate X-chromosome palindrome arms compared to flanking sequence

Previous studies have proposed that high rates of gene conversion in sex chromosome palindromes could lead to elevated GC content in palindrome arms (Caceres et al. 2007, Hallast et al. 2013). We calculated GC content for primate X-chromosome palindrome arms relative to flanking sequence, and found significantly higher median GC content in palindrome arms than in flanking sequence across all three species: 46.3% versus 41.2% (human), 46.3% versus 40.9% (chimpanzee), and 45.2% versus 41.0% (rhesus macaque) (p<0.05 for all three species, Mann-Whitney U) (Figure 1A). The GC content of flanking sequences is slightly elevated compared to the overall GC content of the human X chromosome (39.5%), while the GC content of palindrome arms is markedly higher. The trend of elevated GC content in palindrome arms was highly consistent across different palindromes, with at least eleven out of twelve palindromes having significantly higher GC content in palindrome arms than flanking sequence within each species (p<1 × 10^−6^ for each significant palindrome, chi-square test with Yates correction, Supplemental Table 1). Given that ten out of twelve conserved primate X-chromosome palindrome arms contain one or more protein-coding genes (Jackson et al. 2020), which tend to be GC-rich, we considered the possibility that elevated GC content in primate X-chromosome palindrome arms results from an enrichment of protein-coding genes. However, the difference between GC content in palindrome arms and flanking sequence remained significant after masking protein-coding genes plus their promoters (defined as 1 kb upstream): 44.1% versus 40.1% (human), 44.2% versus 40.1% (chimpanzee), and 44.1% versus 40.5% (rhesus macaque) (p<0.05 for all three species, Mann-Whitney U) (Figure 1B). We conclude that high gene conversion rates in primate X-chromosome palindrome arms are associated with elevated GC content, consistent with the hypothesis that frequent gene conversion causes an increase in GC content over time.

Previous studies of molecular evolution in sex chromosome palindromes have used two different genomic regions as controls for comparison to palindrome arms: Flanking sequence (Caceres et al. 2007, Swanepoel et al. 2020), or the unique sequence that separates palindrome arms, called the spacer (Rozen et al. 2003, Geraldes et al. 2010, Hallast et al. 2013). Given that both spacers and flanking sequence comprise unique sequence, their GC content might be expected to be similar. However, we found that the GC content of spacers occupied an intermediate range between arms and flanking sequence, and did not differ significantly from palindrome arms (Figure 1A, B). This finding may be explained by a recent observation that palindrome spacers are structurally unstable on the timescale of primate evolution: For 7/12 palindromes conserved between human and rhesus macaque, spacer sequence could not be aligned between species, and for five palindromes, part of the spacer from one species corresponded to arm sequence in the other (Jackson et al. 2020). We suggest that palindrome spacers display an intermediate level of GC content because some spacers spent part of their evolutionary history in the palindrome arm, where they were subject to higher levels of gene conversion. There were also examples of X-chromosome palindromes for which part of the arm in one species corresponded to flanking sequence in another (e.g., P9 in human and rhesus macaque, Jackson et al. 2020); this phenomenon may explain why flanking sequence has slightly higher GC content than the X chromosome average, as noted above.

### Nucleotide replacement patterns in human and chimpanzee X-chromosome palindrome arms demonstrate that GC content has increased in the past seven million years

We next looked for evidence of GC-biased gene conversion based on nucleotide replacement patterns in palindrome arms. For each conserved X-chromosome palindrome, we generated a six-way alignment using both palindrome arms from human, chimpanzee, and rhesus macaque. We then identified nucleotide replacements that occurred in the human lineage by searching for sites with the same base in both human arms (e.g. G/G) and a different base in rhesus macaque and chimpanzee arms (e.g. A/A in both species) (Figure 2A). Such fixed differences can be inferred to have arisen through a substitution in the human lineage, followed by gene conversion between human arms (Hallast et al. 2013, Supplemental Note 1). We compared the base composition of the ancestral base at each site of inferred gene conversion to the derived base. If gene conversion is GC-biased, then derived bases should have a higher GC content than ancestral bases. Indeed, we found that the median GC content of derived bases was 64.5%, compared to 41.5% for ancestral bases (p<0.0001, Mann-Whitney U) (Figure 2B). We repeated the same analysis for nucleotide replacements in the chimpanzee lineage, with similar results (62.7% vs 39.4%, p<0.0001, Mann-Whitney U) (Figure 2B). In contrast, a comparable analysis examining the GC content of ancestral versus derived sequence for flanking sequence, using three-way alignments between species, revealed little or no significant difference in base-pair composition (Figure 2C). We conclude that GC-biased gene conversion in human and chimpanzee palindrome arms over the past 7 million years has skewed nucleotide replacement patterns, resulting in derived bases being more than one-and-a-half times more GC rich than the ancestral bases that they replaced.

### Simulations of palindrome gene conversion are consistent with GC bias of about 0.7

Our interpretation of the results shown in Figure 2B assumes that all nucleotide replacements result from the same series of evolutionary events, i.e., a substitution followed by gene conversion. Although we consider this the most parsimonious explanation for fixed differences found in a single species, other explanations cannot be excluded (see Supplemental Note 1). We therefore devised a series of Markov chain Monte Carlo (MCMC) simulations to model palindrome evolution under different magnitudes of GC-biased gene conversion. These simulations allowed us to examine the expected behaviors of palindrome evolution within reasonable parameters for substitution rate, neutral substitution patterns, gene conversion rate, and the magnitude of GC bias, without requiring assumptions about the specific evolutionary trajectory of each site. Our simulations were designed to achieve three objectives: 1) determine the likelihood of observing the pattern of nucleotide replacements shown in Figure 2B in the absence of GC-biased gene conversion, 2) find the magnitude of GC-biased gene conversion most consistent with our results in Figure 2B, and 3) determine what fraction of the elevated GC content seen in primate X-chromosome palindrome arms relative to flanking sequence can be attributed to GC-biased gene conversion. While the simulations shown in Figure 3 were run using identical evolutionary parameters except for the magnitude of GC bias, the effects of altering other parameters are explored in Supplemental Notes 2 and 3; none of these parameter modifications altered the major conclusions of these analyses.

Our simulations model the evolution of a palindrome that was present in the common ancestor of human, chimpanzee, and rhesus macaque, and maintained in all three lineages over 29 million years until the present (see Methods). Briefly, for each iteration, we initialized an ancestral palindrome conforming to the median characteristics of twelve conserved primate X palindromes, including arm length, total GC content, and arm-to-arm identity. We then subjected the ancestral palindrome to rounds of nucleotide substitution followed by gene conversion, with each round representing one generation (Figure 3A). We determined neutral substitution patterns based on alignments of 3.8 Mb gene-masked flanking sequence; our observed pattern showed a strong preference for transitions over transversions, as well as a preference for GC→AT substitutions over AT→GC substitutions, consistent with previous reports (Petrov and Hartl 1999, Zhang and Gerstein 2003, Duret and Arndt 2008; see Methods). We included two branching events to account for the divergence of each lineage, resulting in three evolved palindromes representing those present today in human, chimpanzee, and rhesus macaque. Each simulation described below represents one hundred trials, each simulating twelve independent palindromes, representative of the twelve palindromes described in Figures 1 and 2.

We first used our simulations to determine the likelihood of observing a median difference in GC content between ancestral bases and derived bases as large as that observed in Figure 2B in the absence of GC-biased gene conversion (GC bias = 0.50). For simplicity, we report only the results of evolved human palindromes, given that the palindromes designated as “human” and “chimpanzee” underwent equivalent evolutionary trajectories in our simulations. Out of 100 simulations run without GC-biased gene conversion, we never observed a median difference in GC content between ancestral and derived bases as large as the true median difference of ∼23% in primate X-chromosome palindromes (Figure 3B,C, Figure 2B). Indeed, all observed differences were less than zero, demonstrating that in the absence of GC bias, ancestral bases are expected to be more GC-rich than derived bases, reflecting the higher rate of GC→AT substitutions versus AT→GC substitutions (Figure 3B, C). We conclude that our observed pattern of nucleotide replacements in Figure 2B is unlikely (p<0.01, bootstrapping) in the absence of GC-biased gene conversion.

We next asked what magnitude of GC-biased gene conversion could best explain our observed results in Figure 2B. We repeated our simulations using magnitudes of GC bias ranging from 0.60 to 0.80. Simulations using GC bias of 0.75 and 0.80 both produced median differences in GC content between ancestral and derived bases that were significantly larger than our observed value of 23% (39.0% and 31.8%, respectively, p<0.01 for both), while simulations using GC bias of 0.60 and 0.65 produced values that were significantly smaller (6.8% and 13.8%, respectively, p<.01 and p<0.01) (Figure 3C). We found that an intermediate value of 0.70 produced results highly consistent with our observations, with a median difference in GC content between ancestral and derived bases of 21.8% (ns, Figure 3C). We conclude that our results in Figure 2B are best explained by a magnitude of GC bias of approximately 0.70, consistent with previous estimates derived from analyses of human meioses (Williams et al. 2015, Halldorsson et al. 2016).

Finally, we used our simulations to explore the increase in GC content in palindrome arms that would be produced by GC-biased gene conversion of our inferred magnitude, 0.70, over 29 million years of evolution. In particular, we asked what fraction of the difference in GC content observed between palindrome arms and flanking sequence—ranging from 3.6% in rhesus macaque to 4.1% in chimpanzee, after masking protein-coding genes (Figure 1)—could be explained by GC-biased gene conversion over this time scale. We compared the GC content in simulated human, chimpanzee, and rhesus macaque arms to the GC content of the ancestral palindrome. While the three evolved palindromes had significantly higher GC content than the ancestral palindrome, it was by a median magnitude of 0.68%, explaining at most 19% of our observed difference from primate X-chromosome palindromes (Figure 3D). While GC-biased gene conversion leads to a significant increase in GC content over time, our results suggest that an increase of the magnitude we observed in Figure 1 is unlikely to have occurred since the divergence of human, chimpanzee and rhesus macaque. We conclude that either primate X-chromosome palindromes are considerably older than 29 million years, or that other factors contribute to the difference (see Discussion).

## DISCUSSION

GC-biased gene conversion is a powerful force that shapes nucleotide composition across mammalian genomes (Galtier et al. 2001, Marais 2003, Duret and Galtier 2009). Previous reports have estimated the magnitude of GC bias in humans to be around 68%, based on the detection of autosomal gene conversion events from three-generation pedigrees (Williams et al. 2015, Halldorsson et al. 2016). Here, we inferred a magnitude of GC bias of around 70% in a unique system of twelve large palindromes conserved on the X chromosome, using a comparative genomic approach combined with evolutionary simulations. The concordance between our results and those of previous studies, including investigations of GC-biased gene conversion in human Y chromosome palindromes (Hallast et al. 2013, Skov et al. 2017), suggests that the magnitude of GC bias in humans is relatively constant across diverse genomic contexts. From this, we further infer that regional differences in the effects of GC-biased gene conversion—such as the GC-skewed nucleotide replacements that we detect in primate X-chromosome palindrome arms—stem from regional differences in the rate of gene conversion, rather than in the strength of GC bias.

Despite the prediction that high rates of gene conversion could amplify the effects of GC-biased gene conversion, few previous studies have examined the GC content of sex chromosome palindrome arms. One human X chromosome palindrome with putative orthologs in other mammals was found to have higher GC content in palindrome arms compared to flanking sequence in all sixteen species studied (Caceres et al. 2007). Results based on six human Y chromosome palindromes were mixed, with two palindromes showing significantly higher GC content in arms than in spacer, and the other four palindromes showing no significant difference (Hallast et al. 2013). The selection of the spacer for comparison may have reduced the significance of the latter findings, given that we found significant results only from comparing GC content between palindrome arms and flanking sequence. In general, we propose that flanking sequence represents a stronger comparison than spacers for molecular analyses of palindrome evolution, due to the fact some X-chromosome palindrome spacers have mixed evolutionary histories that may include time spent within the palindrome arm (Jackson et al. 2020).

Although we found that GC content in primate X-chromosome palindromes is robustly elevated in palindrome arms versus flanking sequence, simulations show that less than 20% of this increase can be attributed to GC-biased gene conversion since the divergence of the human and rhesus macaque lineages. One possible explanation is that palindromes arose much earlier in primate or mammalian evolution, resulting in additional time to accumulate GC content. However, given the order-of-magnitude difference between our observed results and simulations, we consider under-estimation of palindrome age unlikely to explain the entire discrepancy. We instead propose two mutually compatible possibilities: that GC-rich sequence is more susceptible to palindrome formation, and/or that GC-rich palindromes are more likely to survive over long evolutionary timescales. Both possibilities are bolstered by the fact that although high rates of recombination can elevate GC content over time (Montoya-Burgos et al. 2003, Li et al. 2016), elevated GC content can also increase local rates of recombination (Petes and Merker 2002, Kiktev et al. 2018). Given that palindrome formation is believed to require two recombination events (Kuroda-Kawaguchi et al. 2001), recombinogenic GC-rich sequence may be more likely than AT-rich sequence to form palindromes. Palindromes with high GC content may also have a survival advantage over palindromes with lower GC content, given that high rates of recombination are required to prevent arms from diverging over time. We speculate that both factors—an increased tendency for GC-rich sequence to form and maintain palindromes, combined with further gains in GC content over time from GC-biased gene conversion— contribute to the remarkably GC-rich palindromes we observe in X-chromosome palindromes from human, chimpanzee and rhesus macaque.

## DATA AVAILABILITY

BAC sequences used for this study are available from GenBank (https://www.ncbi.nlm.nih.gov/) under accession numbers listed in Supplemental Table 2. The authors affirm that all other data necessary for confirming the conclusions of the article are present within the article, figures and tables. Code used to generate the simulated data can be found at https://github.com/ejackson054/GC-biased-gene-conversion.

## ACKNOWLEDGEMENTS

This work was supported by the Howard Hughes Medical Institute, the Whitehead Institute, and generous gifts from Brit and Alexander d’Arbeloff, Arthur W. and Carol Tobin Brill, and Matthew Brill.

---

**Supplemental Table 1:** GC content is elevated in primate X palindrome arms relative to flanking sequence. P-values are from chi-square test with Yates correction.

| Palindrome | Arm 1 length | Flanking length | Arm 1 GC | Flanking GC | p-value |
| --- | --- | --- | --- | --- | --- |
| P2 | 24849 | 83517 | 44.07 | 39.22 | 1.80E-266 |
| P3 | 36398 | 83392 | 47.08 | 37.52 | 0 |
| P6 | 37939 | 203740 | 48.05 | 43.91 | 0 |
| P7 | 26581 | 140783 | 43.69 | 38.27 | 0 |
| P8 | 57336 | 56225 | 45.86 | 47.62 | 3.50E-07 |
| P9 | 119125 | 92765 | 43.19 | 40.21 | 9.95E-92 |
| P10 | 9193 | 101002 | 50.36 | 42.21 | 1.34E-142 |
| P11 | 140582 | 176247 | 41.72 | 39.47 | 0 |
| P21 | 46811 | 88685 | 46.83 | 42.73 | 8.19E-160 |
| P24 | 10050 | 92241 | 61.11 | 50.83 | 3.14E-138 |
| P25 | 35448 | 288582 | 53.01 | 48.46 | 3.76E-310 |
| P26 | 50037 | 250270 | 41.54 | 37.61 | 7.62E-155 |

Chimpanzee
| Palindrome | Arm 1 length | Flanking length | Arm 1 GC | Flanking GC | p-value |
| --- | --- | --- | --- | --- | --- |
| P2 | 25099 | 82455 | 43.95 | 39.36 | 2.25E-257 |
| P3 | 36247 | 83212 | 47.1 | 37.53 | 0 |
| P6 | 34932 | 196227 | 48.89 | 44.14 | 0 |
| P7 | 28529 | 140226 | 44.2 | 38.26 | 0 |
| P8 | 53145 | 65431 | 45.94 | 47.86 | 1.05E-14 |
| P9 | 119105 | 88148 | 43.08 | 39.74 | 1.07E-116 |
| P10 | 7591 | 100457 | 46.62 | 42.05 | 2.27E-163 |
| P11 | 160191 | 185640 | 41.63 | 39.19 | 0 |
| P21 | 36887 | 94054 | 46.6 | 42.53 | 6.08E-227 |
| P24 | 10712 | 90734 | 59.62 | 50.86 | 2.49E-63 |
| P25 | 34937 | 289807 | 52.74 | 47.95 | 2.79E-303 |
| P26 | 46893 | 240223 | 41.2 | 37.91 | 2.17E-138 |

Rhesus macaque
| Palindrome | Arm 1 length | Flanking length | Arm 1 GC | Flanking GC | p-value |
| --- | --- | --- | --- | --- | --- |
| P2 | 42623 | 91615 | 44.32 | 39.06 | 0 |
| P3 | 34782 | 96783 | 47.07 | 37.85 | 0 |
| P6 | 15564 | 221983 | 48.34 | 44.79 | 0 |
| P7 | 24152 | 187629 | 43.93 | 38.3 | 0 |
| P8 | 38447 | 89961 | 47.77 | 45.91 | 0 |
| P9 | 81966 | 152376 | 43.61 | 40.54 | 0 |
| P10 | 6561 | 117616 | 46.14 | 41.55 | 1.65E-172 |
| P11 | 105994 | 188318 | 42.46 | 39.55 | 0 |
| P21 | 21963 | 89617 | 44.35 | 42.31 | 8.97E-267 |
| P24 | 7673 | 90082 | 61.05 | 50.28 | 0 |
| P25 | 101313 | 273906 | 51.23 | 47.45 | 0 |
| P26 | 45037 | 198305 | 41.74 | 37.11 | 0 |

**Supplemental Table 2:** GenBank accession numbers for chimpanzee and rhesus macaque clones analyzed for this project

| Clone | Accession | Palindrome |
| --- | --- | --- |
| CH250-106M20 | AC280444 | P7 |
| CH250-114J18 | AC280531 | P9/P10 |
| CH250-136N6 | AC280430 | P3 |
| CH250-137I15 | AC280580 | P26 |
| CH250-168E3 | AC280520 | P21 |
| CH250-174F12 | AC280455 | P21 |
| CH250-191K20 | AC280440 | P2 |
| CH250-197O3 | AC280457 | P6 |
| CH250-228D11 | AC280538 | P11 |
| CH250-236O7 | AC280541 | P8 |
| CH250-371L16 | AC280564 | P9 |
| CH250-398K19 | AC280504 | P21 |
| CH250-412K19 | AC280483 | P6 |
| CH250-417G7 | AC280442 | P26 |
| CH250-424H13 | AC280467 | P25 |
| CH250-462M8 | AC280473 | P21 |
| CH250-487N16 | AC280453 | P26 |
| CH250-491H11 | AC280464 | P24 |
| CH250-493M11 | AC280503 | P25 |
| CH250-498I16 | AC280468 | P11 |
| CH250-503C21 | AC280489 | P25 |
| CH250-503N19 | AC280524 | P25 |
| CH250-540J3 | AC280456 | P11 |
| CH250-541H5 | AC280425 | P3 |
| CH250-563M7 | AC280492 | P3 |
| CH250-80G22 | AC280568 | P9 |
| CH250-87B7 | AC280549 | P7 |
| CH250-94G2 | AC280518 | P3 |
| CH251-161L14 | AC280465 | P7 |
| CH251-177B21 | AC280525 | P2 |
| CH251-183G21 | AC280578 | P8 |
| CH251-239P10 | AC280533 | P26 |
| CH251-240O17 | AC280544 | P6 |
| CH251-277H18 | AC280556 | P7 |
| CH251-285D14 | AC280499 | P26 |
| CH251-346A10 | AC280446 | P25 |
| CH251-397P16 | AC280507 | P3 |
| CH251-4M24 | AC280567 | P11 |
| CH251-504H5 | AC280540 | P10 |
| CH251-506D4 | AC280415 | P6 |
| CH251-514B7 | AC280488 | P8 |
| CH251-542A6 | AC280462 | P10 |
| CH251-565G15 | AC280459 | P21 |
| CH251-635P13 | AC280558 | P11 |
| CH251-651H9 | AC280522 | P11 |
| CH251-657L4 | AC280546 | P9/P10 |
| CH251-658J15 | AC280445 | P8 |
| CH251-65E21 | AC280530 | P24 |
| CH251-671I19 | AC280576 | P3 |
| CH251-677L24 | AC280565 | P24/P25 |
| CH251-737G9 | AC280491 | P9/P10 |
| CH251-73C22 | AC280423 | P2 |

**Supplemental Note 1:** Inference of gene conversion from conserved palindromes in human, chimpanzee, and rhesus macaque

Using the logic of Hallast et al. 2013, we inferred gene conversion events from fixed nucleotide replacements in X palindrome arms that occurred in either human or chimpanzee, using rhesus macaque as an outgroup. For simplicity, the scenarios below describe a fixed replacement in the human lineage. We propose that fixed nucleotide replacements result from a substitution in humans after divergence from chimpanzee, followed by gene conversion that homogenizes the substitution between arms (Scenario 1). In theory, other scenarios could lead to the same result. In one alternative scenario, the ancestral palindrome was heterozygous at the site in question, with gene conversion occurring in one direction in rhesus macaque and chimpanzee, and the opposite direction in human (Scenario 2). We consider this scenario highly unlikely because it requires the initial site to remain heterozygous for 1.1 million generations before undergoing gene conversion in human and chimpanzee (see Figure 3A). Given our inferred intrachromosomal gene conversion rate of 4.5 × 10^−5^ events per nucleotide per generation, the probability of any given site not undergoing gene conversion over 1.1 million generations is (1 - 4.5 × 10^−5^) ^ 1100000, which is effectively zero (<2.22 × 10^−308^).

We also considered a scenario in which the initial substitution occurred in the human-chimpanzee common ancestor, then underwent gene conversion in opposite directions in human and chimpanzee (Scenario 3). Given that X palindromes have on average only 1 difference between arms for every 2200 nucleotides, this scenario could explain at most observed nucleotide replacements in 1 out of 2200 positions in X palindrome arms (0.045%), if all heterozygous sites resolved in opposite directions in each lineage. We observed nucleotide replacements in 2567 out of 409,579 positions in X palindrome arms (0.65%), suggesting that Scenario 3 can account for no more than 7% (0.045% / 0.65%) of our observations.

Finally, we used evolutionary simulations with event tracing to estimate what fraction of fixed nucleotide replacements in human, chimpanzee, and rhesus macaque would arise through each scenario under reasonable evolutionary parameters (see Figure 3, Methods). We found that the vast majority of fixed nucleotide replacements (93.3%) arose through Scenario 1, while around 2.5% arose through Scenario 3. As predicted, we never observed fixed replacements arising from Scenario 2. The remaining fixed nucleotide replacements (4.2%) resulted from other scenarios that involved multiple substitution events. Importantly, our conclusions in Figure 3 are agnostic to the method by which each fixed nucleotide replacement arose, and depend only on the ability of a given set of evolutionary parameters to reproduce the replacement patterns seen in Figure 2B.

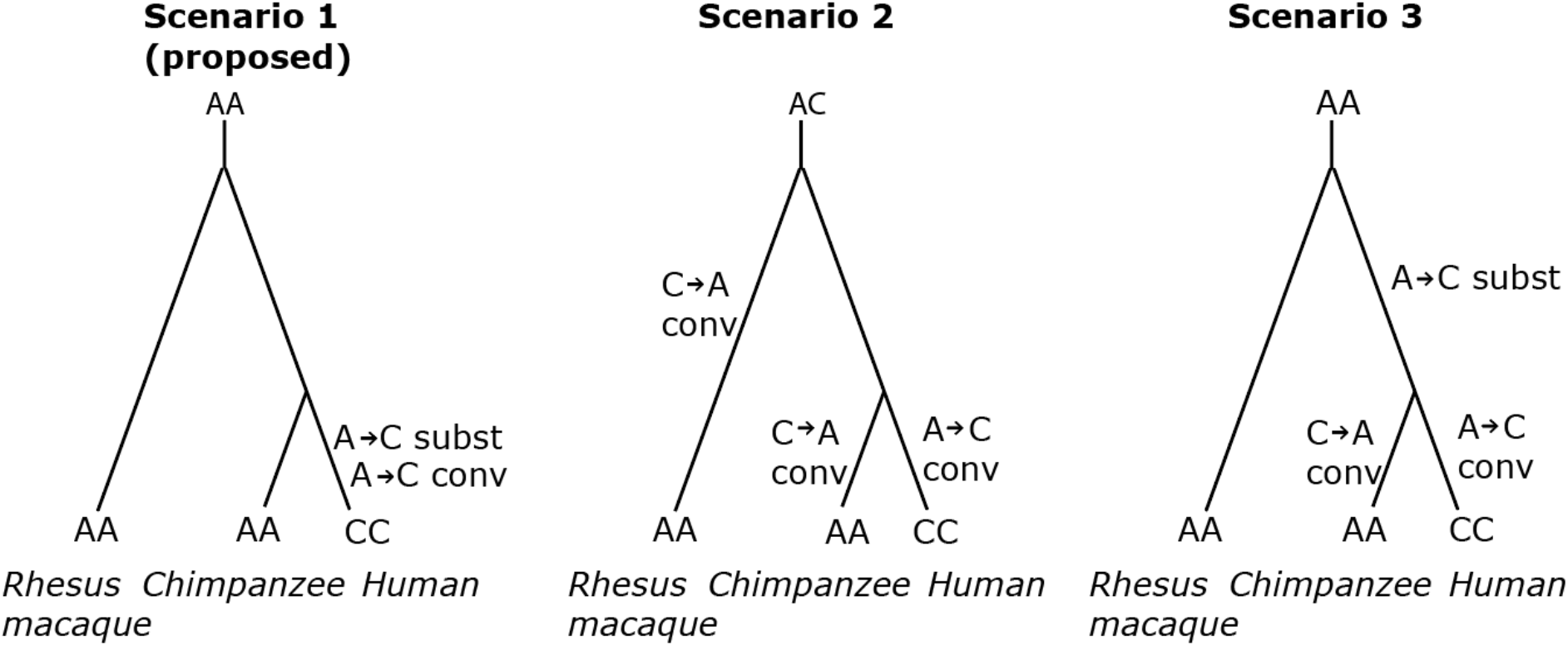

**Supplemental Note 2:** Effects of altering substitution rates and generation times for simulations

Although our calculations of generation numbers and substitution rates for each branch of the evolutionary tree were based on estimates of divergence times and divergence times from recent literature, these values nevertheless are subject to uncertainty. For example, while we estimated a substitution rate of 1.20 × 10^−8^ substitutions per nucleotide per generation for Branches 3 and 4 based on observed divergence of 0.84% between human and chimpanzee and assuming 700,000 generations, the observed divergence between human and chimpanzee could instead have resulted from a higher substitution rate combined with a lower number of generations, or vice versa.

To test the effect of this uncertainty on our simulations, we considered two fairly extreme cases: 1) The true substitution rates in all lineages were twice as high as we estimated, while the true number of generations was halved; 2) The true substitution rates in all lineages were half as high as we estimated, while the true number of generations was doubled. We then repeated our simulations for both of these cases (“high substitution” and “low substitution”) and compared our results to the results obtained using our original calculations. We find that these alterations make no difference to our inference of the magnitude of GC bias at 0.70 (difference between observed and simulated results not significant, p>0.05).

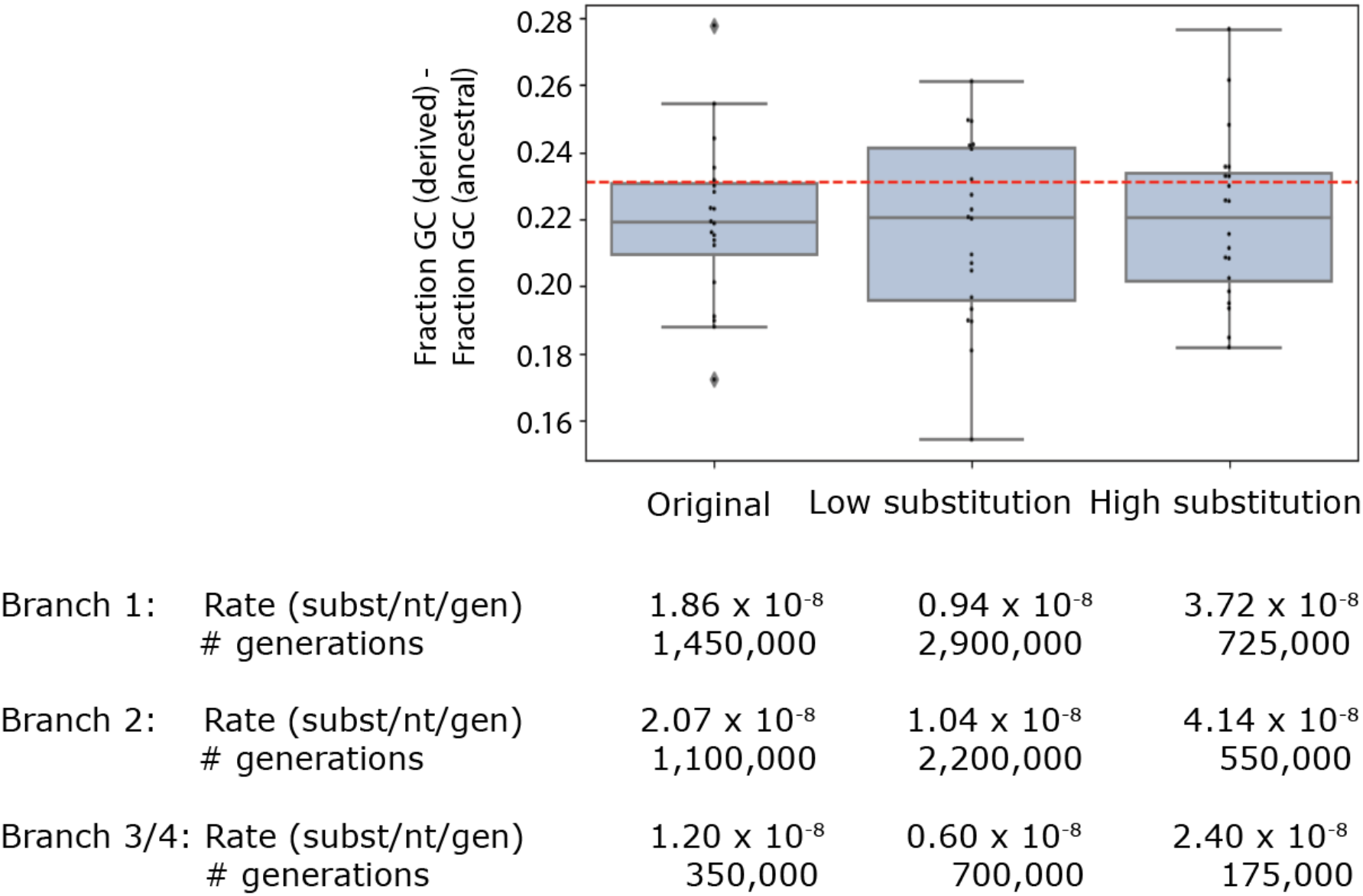

**Supplemental Note 3:** Effects of altering the neutral substitution matrix for simulations

We considered the effect that altering the neutral substitution matrix would have on our simulations. In particular, we considered two possible limitations of our inferred neutral substitution matrix. First, our matrix was derived from flanking X chromosome sequence with a median GC content around 40%, while palindrome arms have a median GC content around 46%. Previous work has shown that substitution patterns can differ based on regional GC content, with regions with high GC content showing a lower rate of strong (GC) to weak (AT) mutations (Duret and Arndt 2008). Using a matrix derived from sequence with a lower GC content could in theory lead to over-estimation of AT mutation bias, and subsequent over-estimation of the GC conversion bias required to balance it. We therefore re-calculated our neutral substitution matrix using a subset of flanking sequence (1.3 Mb) with a total GC content of 45% and repeated our simulations (figure below, top panel). Our inference of GC bias remained unchanged: Using 20 simulations of 12 palindromes each, our observed results were still most consistent with a GC bias magnitude of 0.70 (difference between observed and simulated results not significant, p>0.05).

The second limitation we considered was our use of parsimony to infer substitution events for our neutral substitution matrix. While this is appropriate for most types of substitutions on the time scale of human-chimpanzee evolution, it has been shown to under-estimate the frequency of CpG substitutions, which occur more frequently than other substitutions and can thus occur twice at the same site (Duret 2006). Under-estimation of CpG substitutions could lead us to under-estimate the AT mutation bias, and therefore under-estimate the magnitude of GC conversion bias. To determine the impact on our simulations, we re-estimated our rate of CpG substitutions to align with the values found by Duret and Arndt 2008, who found that CpG substitutions (CG→AT at CpG sites) are 14 times more common than the same substitutions at non-CpG sites using a maximum-likelihood method. While adjusting the frequency of CpG adjustments shifted our simulated differences in GC content between derived and ancestral bases slightly downwards, our results were still consistent with a magnitude of GC bias if 0.70 (difference between observed and simulated results not significant, p>0.05) We conclude that our results are robust to reasonable shifts in the neutral substitution matrix.

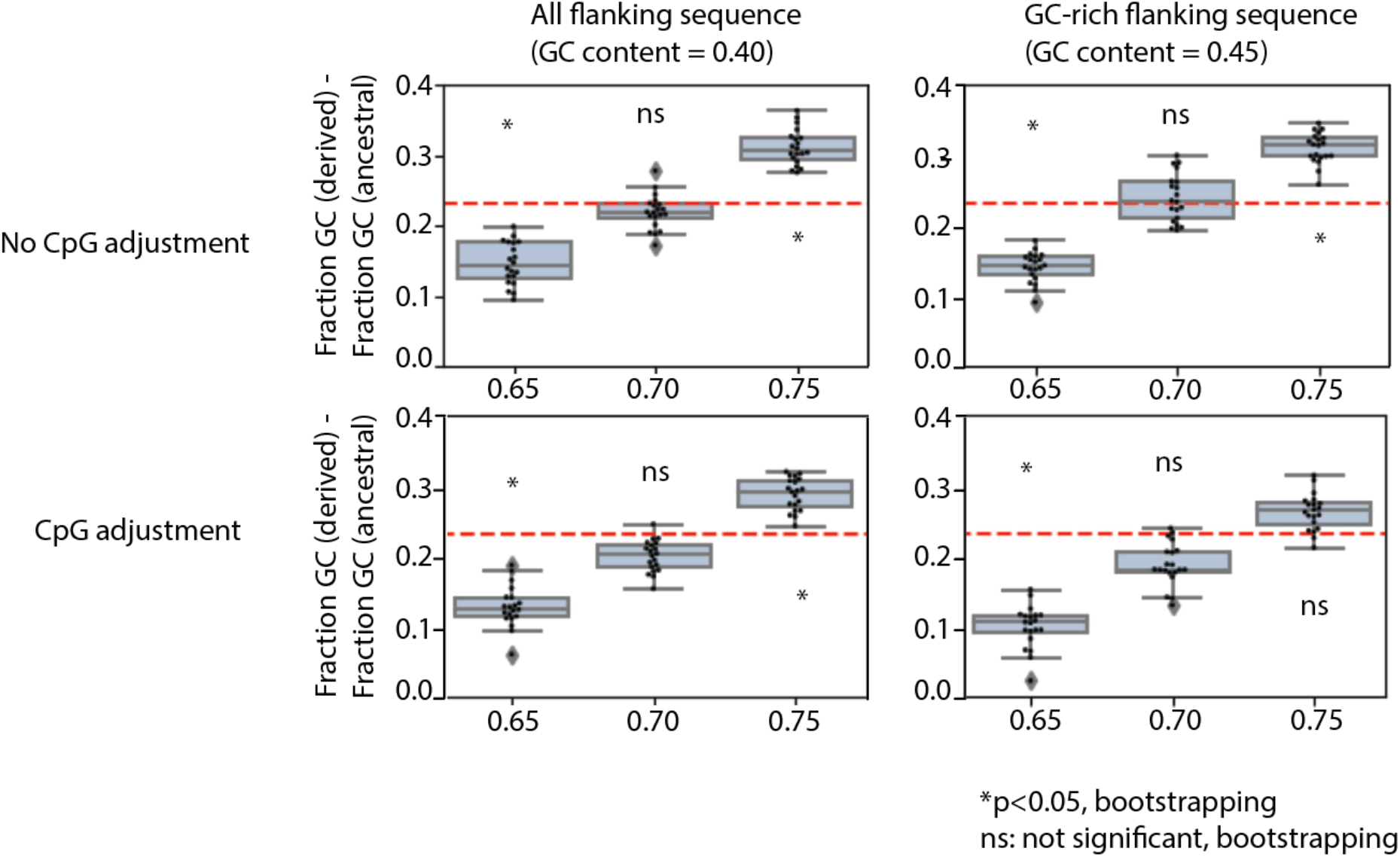

